# A Global Transcriptomic Analysis of *Staphylococcus aureus* Biofilm Formation Across Diverse Clonal Lineages

**DOI:** 10.1101/2020.12.17.423160

**Authors:** Brooke R. Tomlinson, Morgan E. Malof, Lindsey N. Shaw

## Abstract

A key characteristic of *S. aureus* infections, and one that also varies phenotypically between clones, is that of biofilm formation, which aids in bacterial persistence through increased adherence and immune evasion. Though there is a general understanding of the process of biofilm formation - adhesion, proliferation, maturation, and dispersal - the tightly orchestrated molecular events behind each stage, and what drives variation between *S. aureus* strains, has yet to be unraveled. Herein we measure biofilm progression and dispersal in real-time across the 5 major *S. aureus* CDC-types (USA100-USA500) revealing adherence patterns that differ markedly amongst strains. To gain insight into this, we performed transcriptomic profiling on these isolates at multiple time points, compared to planktonically growing counterparts. Our findings support a model in which eDNA release, followed by increased positive surface charge, perhaps drives initial abiotic attachment. This is seemingly followed by cooperative repression of autolysis and activation of PNAG production, which may indicate a developmental shift in structuring the biofilm matrix. As biofilms mature, diminished translational capacity was apparent, with 53% of all ribosomal proteins downregulated, followed by upregulation of anaerobic respiration enzymes. These findings are noteworthy because reduced cellular activity and an altered metabolic state have been previously shown to contribute to higher antibiotic tolerance and bacterial persistence. In sum, this work is, to our knowledge, the first study to investigate transcriptional regulation during the early, establishing phase of biofilm formation, and to compare global transcriptional regulation both temporally and across multiple clonal lineages.

**Importance:** *Staphylococcus aureus* is a highly virulent, opportunistic pathogen and a leading cause of both nosocomial and community acquired infections. Biofilms are associated with persistent, chronic infections and the capacity to form a biofilm varies phenotypically amongst *S. aureus* clonal lineages. The molecular regulation of biofilm formation has been a popular area of study for this pathogen, however, a comprehensive mapping of the regulatory processes and factors driving strain-specific variation has yet to be elucidated. This study presents transcriptomic analyses of five diverse methicillin-resistant *S. aureus* isolates during various stages of biofilm formation, tracked in real time. The transcriptomic profiles of all five isolates were compared to identify both core and unique networks of regulation. Importantly, much of what we currently know about biofilms is based on mature, preformed biofilm populations, with little known of the transient regulation driving attachment, proliferation, maturation, and dissemination. We address this issue by investigating transcriptional regulation during the early, establishing phase of biofilm formation, and compare global transcriptional regulation both temporally and across multiple clonal lineages. This study provides a launching point towards understanding the highly orchestrated regulation driving each phase of biofilm development and may inform on future strategies to combat biofilm-mediated infections.

**Data Summary:**

1. RNA sequencing results have been deposited to the NCBI Gene Expression Omnibus; GEO submission GSE163153 (url – https://www.ncbi.nlm.nih.gov/geo/query/acc.cgi?acc=GSE163153)
2. Overview and comparisons of gene expression within biofilm and planktonic cell populations for the various *Staphylococcus aureus* strains are shown in Table S1, S2, S3. Validation and visual depiction of these data can be found as Figure S2 and S3 respectively (available in supplementary material).

## Introduction

More than 2.8 million antibiotic resistant infections occur in the U.S. annually (1) and with an imminent post-antibiotic era in our future, aggressive action is needed to develop new treatment and prevention strategies. *Staphylococcus aureus* is a highly virulent, opportunistic pathogen and is responsible for > 320,000 of these reported infections (1). Ubiquitous in hospital settings, *S. aureus* persistently colonizes one third of the population (2, 3) and is a leading cause of indwelling device infections caused by biofilms (4, 5). In general, device-associated infections initiate through the adherence of bacterial cells to the surface of an implant, such as catheters, heart valves, or prosthetic joints. Once attached, bacteria secrete DNA, polysaccharides, and proteins, forming an extracellular polymeric matrix. Within biofilms, there is population heterogeneity, the formation of persister cells, upregulation of stress responses, and altered microclimates (reviewed in (6)). Consequently, a combination of each of these factors impact drug stability/penetration and render *S. aureus* and other biofilm forming pathogens broadly resistant to the majority of our antimicrobial arsenal (7). Biofilms are also associated with persistent, chronic infections and require aggressive treatment tactics, such as implant removal and extensive debriding of infected tissue and bone (8, 9).

When an implant is introduced to the body the immune response readily coats its abiotic surface with host proteins, including fibrinogen, fibronectin and laminin (10, 11). At this point, *S. aureus* begins the highly orchestrated, cyclical process of biofilm formation: attachment, proliferation, maturation, and detachment. This process commences with production of host colonization and intercellular adhesion factors (12), such as <u>m</u>icrobial <u>s</u>urface <u>c</u>omponents <u>r</u>ecognizing <u>a</u>dhesive <u>m</u>atrix <u>m</u>olecules (MSCRAMMs). These include clumping factors (ClfA, ClfB) (13), fibronectin-binding proteins (FnbA, FnbB), and the serine aspartate repeat proteins (SdrC, SdrD, and SdrE) (14), all of which associate with and tightly adhere to host proteins to promote colonization of implant surfaces. Once attached, biofilm proliferation occurs via secreted DNA, polysaccharides, and proteins. The peptidoglycan hydrolase, AtlA, promotes release of these cytosolic factors through autolysis (15), whilst the Ica proteins (IcaADBC) contribute to the matrix by synthesizing polysaccharide intercellular adhesin (PIA). Proliferating cells within this matrix lose direct contact with the implant surface and host proteins, causing them to rely on cell-cell and cell-<u>e</u>xtracellular <u>p</u>olymeric <u>s</u>ubstance (EPS) adhesion (12). Some MSCRAMMS, such as SdrC and FnBPs, can self-associate to serve this role (14, 16), but IgG Binding Protein A (Spa) is the dominant component of the extracellular matrix that facilitates bacterial aggregation (17). As a biofilm matures, microcolony clusters begin to exhibit different growth characteristics and protein expression depending on their location within the biofilm (18). Eventually, the biofilm disperses or is disrupted, shedding these clusters to spread and repeat the process elsewhere.

The capacity to form a biofilm varies amongst *S. aureus* strains, as does their ability to cause disease. Methicillin-resistant *S. aureus* strains are most commonly categorized by the universal generic subtyping method, pulsed-field gel electrophoresis (PFGE). Specific *S. aureus* PFGE clonal groups have been correlated to patient outcomes, outbreaks, and pathogenicity (19–21). For example, PFGE-types USA300 and USA400 are most often associated with community-acquired (CA) infections (22), whereas, USA100, USA200, and USA500 are associated with hospital-acquired (HA) infections (22). In studies surveying isolates present within the United States, USA300 and USA100 were the most prevalent CA and HA isolates respectively (22–25). USA300 is also the primary epidemic lineage in the United States, causing severe necrotizing pneumonia (26, 27) and skin and soft tissue infections (28). The suspected progenitor to USA300, USA500, is slightly less prevalent despite having a similar capacity for virulence (29). Although less common than USA300 and USA100, USA200 and USA400 strains cause severe and lethal disease, even in healthy individuals. USA400, in particular, is associated with cases of severe sepsis (30), whilst USA200 is a leading cause of endocarditis and toxic shock syndrome (31). USA100 has demonstrated proficiency in causing endocarditis as well (20), but this clonal type is perhaps best known for its proclivity to multidrug resistance (22) and hospital onset infections (23, 32). USA300 and USA500 strains are commonly described as poor biofilm formers (33, 34), with USA300 characterized as marginally better than USA500 (34). Conversely, USA100, USA200, and USA400 strains have been shown to form comparatively stronger biofilms *in vitro* (20, 35, 36).

The diversity of disease progression for *S. aureus* strains is a consequence of differential expression of its arsenal of virulence factors. Although biofilm formation does not strictly correlate with severity of disease, a number of the biofilm factors mentioned above also serve as virulence determinants. For example, beyond promoting cell clumping through fibrinogen binding, ClfA also coats cells with coagulase-generated fibrin fibrils, which impedes phagocytosis by host cells (37). Spa, in addition to promoting adhesion, also facilitates immune evasion by binding host immunoglobulin and impeding phagocytosis (38). Despite mounting evidence of virulence factor production and biofilm formation concomitance (39), little is known about the regulatory networks behind the observed diversity in pathogenicity, infection niche, and biofilm capacity of *S. aureus* isolates. Additionally, most, if not all, studies to date have narrowly focused on mature, established biofilms. Herein, we perform a comprehensive transcriptomic analysis of representative strains from the USA100, USA200, USA300, USA400, and USA500 clonal lineages. We compare the global transcriptional profiles of their biofilms to planktonic counterparts, temporally and across strains, to identify unique responses in biofilms over time. As a result, we uncover several biofilm associated regulatory pathways, some of which are universally employed by all *S. aureus* strains, whilst others allude to strain-specific variations and niche specialization.

## Materials and Methods

### Bacterial strains and growth conditions

Methicillin-resistant *S. aureus* strains from different CDC derived USA clonal lineages (Table 1) were cultured in tryptic soy broth (TSB) at 37°C with agitation (250 rpm). *S. aureus* biofilm cultures were generated in microtiter plates as described previously (40), with the following modifications. Briefly, overnight cultures were normalized to an OD_600_ of 5.0 in phosphate-buffered saline (PBS), before 20 μL was added to 180 μL of fresh TSB in 96-well microtiter plates for a final OD_600_ of 0.5.

**Table 1.** Bacterial strains used in this study.

| Strains | Clonal Type* | Origin<sup>†</sup> | Reference |
| --- | --- | --- | --- |
| N315 | USA100 | HA-MRSA, pharyngeal smear, Japan | (1, 2) |
| MRSA252 | USA200 | HA-MRSA, postoperative infection, UK | (3) |
| LAC | USA300 | CA-MRSA, skin/soft tissue, USA | (4) |
| MW2 | USA400 | CA-MRSA, septicemia/septic arthritis, USA | (5) |
| NRS385 | USA500 | HA-MRSA, bloodstream, USA | (2) |

\*clonal type of each strain as determined by PFGE in previous studies
<sup>†</sup>presentation of infection: HA, nosocomial; CA, community-acquired

### Crystal violet biofilm assays

Following 24 hours of static growth at 37°C, biofilms were washed 3 times with 200 μL PBS and fixed with 100 μL of 100% ethanol. After drying, 200 μL of crystal violet was added, incubated at room temperature for 15 minutes, and aspirated before being washed 3 times with PBS to remove unabsorbed stain. Following a second drying step, 200 μL of 100% ethanol was added to solubilize the crystal violet. Absorbance of the solubilized crystal violet was measured at OD_550_ following 1:10 dilution in PBS. Crystal violet assays were performed in biological triplicate with 8 technical replicates.

### Biofilm formation measurement in real time

A <u>r</u>eal <u>t</u>ime <u>c</u>ell <u>a</u>nalyzer (RTCA) xCELLigence MP (ACEA Bioscience) instrument was used to monitor biofilm formation over time. The RTCA measures adherence of cells based on impedance of electrical signals in dedicated 96-well plates containing electrodes (E-plates), and is expressed as a Cell Index (CI). CI is a relative unit defined as the difference in electrical signal impedance before and after the addition of cells, over time. For analysis, the RTCA was placed in a 37°C incubator for one hour prior to experimentation to allow the instrument temperature to equilibrate. Next, 96-well E-plates were loaded with 180 μL of TSB, positioned in the RTCA, and measured for background signal. Using the same plate, *S. aureus* biofilms were prepared as described above, and statically incubated in the RTCA, with reads taken every 15 minutes for 25 h. The data generated herein is from nine biological replicates per strain.

### RNA sequencing

*S. aureus* biofilms from the various USA pulse field lineages were allowed to form in 96-well microtiter plates as described above and grown in biological triplicate for 5 h, 10 h and 24 h at 37°C in a static incubator. To collect planktonic samples, 75 μL of supernatant was removed from the top of each well and those for like strains were pooled. Samples were immediately combined with 5mL of ice-cold PBS, and pelleted by refrigerated centrifugation. For biofilm samples, the remaining supernatant was removed and biofilm containing wells were washed three times with 200 μL of ice-cold PBS. Ice-cold PBS was added a final time, pipetted vigorously to disrupt biofilm cells, and like strains were pooled. Samples were then immediately combined with an additional 5 mL of ice-cold PBS and pelleted by refrigerated centrifugation. Total RNA was isolated from cell pellets as described previously (41) using an RNeasy Kit (Qiagen) with DNA removed using a TURBO DNA-free kit (Ambion). DNA removal was confirmed by PCR using primers OL398 and OL399 (Table 2) and RNA quality was assessed using an Agilent 2100 Bioanalyzer system with corresponding RNA 6000 Nano kit (Agilent) to confirm RNA integrity (RIN). Samples with a RIN of >9.7 were used in this study. Biological triplicate samples for each strain were then pooled at equal RNA concentrations and ribosomal RNA was removed using a Ribo-Zero Kit for Gram Positive Bacteria (Illumina). Following this, a second round of mRNA enrichment was performed using a MICROBExpress Bacterial mRNA enrichment kit (Agilent). Removal efficiency of rRNA was confirmed using an Agilent 2100 Bioanalyzer system and RNA 6000 Nano kit (Agilent). Enriched mRNA samples were then used for RNA sequencing using an Illumina NextSeq. Library preparation and RNA sequencing was performed following Truseq Stranded mRNA Kit (Illumina) recommendations with mRNA enrichment steps omitted. Quality, concentration, and average fragment size of each sample was assessed using an Agilent 2100 Bioanalyzer system and RNA 6000 Nano kit (Agilent) prior to sequencing. Library concentrations for pooling of barcoded samples was assessed by RT-qPCR with a KAPA Library Quantification kit (KAPA Biosystems) as recommended for high sensitivity. Samples were run on an Illumina NextSeq with a corresponding 150-cycle NextSeq Mid Output Kit v2.5. Experimental data from this study were deposited in the NCBI Gene Expression Omnibus (GEO) database (GEO accession number GSE163153).

**Table 2.**
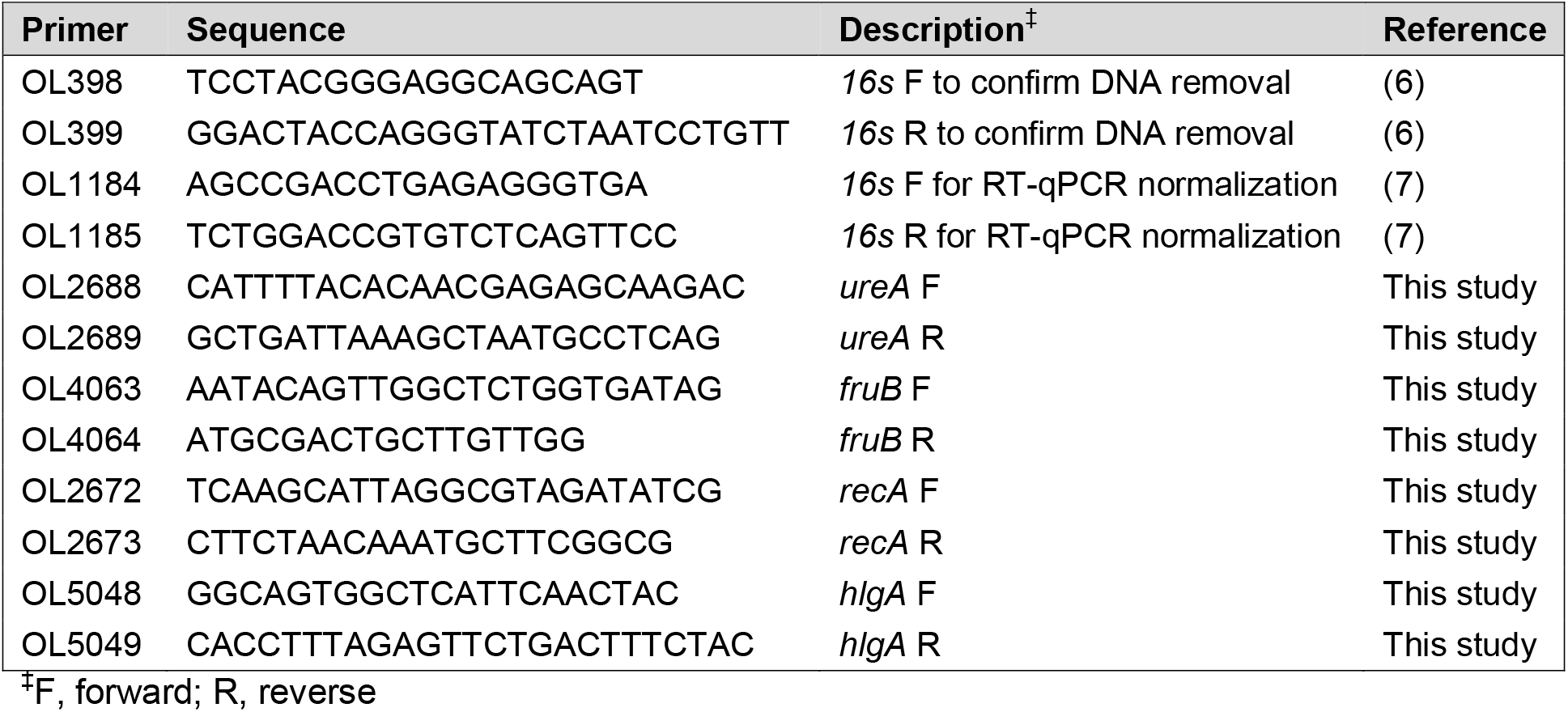
Primers used in this study.

### RNA-seq bioinformatics

Data was exported from BaseSpace (Illumina) in fastq format and analyzed using CLC Genomics Workbench 12 (Qiagen Bioinformatics). Reads were aligned using the RNA-seq Analysis tool to the following reference genomes: USA100 (NC_002745), USA200 (NC_002952), USA300 (NC_007793), USA400 (NC_003923). A well annotated reference genome is not currently available for USA500. USA500 is the suspected progenitor of USA300 (29, 42), therefore, the USA300 reference genome was used. Gene expression values were calculated using the Expression Browser tool and differential expression between biofilm and planktonic samples were generated for each strain and timepoint individually using the Differential Expression in Two Groups tool.

### Identification of homologous genes for intraspecies comparisons

For comparisons across clonal lineages, homologous genes for each reference genome were determined by reciprocal best BLASTn hit. Gene tracks were generated from the respective reference genomes and gene annotations were extracted from each gene track. Extracted annotations were then used as a query for BLASTn against the USA300 genome annotation as a reference and vice versa. Results from the reciprocal BLASTn were then sorted by lowest E-value and those returning in a reciprocal best match were deemed homologous.

### RT-qPCR verification of RNA-seq findings

To validate RNA-seq findings, a random selection of genes were assayed by Real-Time Quantitative Reverse Transcription PCR (RT-qPCR). Strains were grown and harvested as described above for RNA-seq studies. Total RNA was isolated from cell pellets and DNA removal was performed as described above. Samples were reverse transcribed using an iScript cDNA Synthesis Kit (BioRad). RT-qPCR was then performed using gene-specific primers (Table 2) and TB Green Premix Ex Taq (Takara). Levels of gene expression were normalized to that of 16S rRNA and expression was assessed for each biofilm and each timepoint relative to its planktonic counterpart using the 2^−ΔΔCt^ method (43).

## Results and Discussion

### Comparing biofilm formation of Methicillin-resistant *S. aureus* isolates

The *S. aureus* strains used in this study were originally isolated from diverse infection niches and geographic locations (Table 1). To compare their abilities to form a biofilm, we performed classical crystal violet staining assays on mature 24 h biofilms grown in TSB (Fig. 1). Obvious differences were observed in total mass of biofilms at this timepoint, with the USA500 representative having the highest biofilm forming capacity, followed by USA300; with USA100, USA200, and USA400 all displaying similar phenotypes. To explore and compare biofilm formation by these strains more fully, we next employed a more quantitative approach using a Real Time Cell Analyzer (RTCA) device. The RTCA measures adherence of cells based on impedance of electrical signals between electrodes lining each well of a specialized 96-well plate, and expresses these measurements in a relative unit, known as a Cell Index (CI). This allows measurements to be taken at multiple time points on the same biofilm population and forgoes disruptive processing steps. When we performed these studies, we noted that CI values after 24 hours of growth largely correlated to crystal violet assay results, with USA500 and USA300 producing more robust biofilms, and USA100 and USA200 seemingly producing less biomass (Fig. 2). Of note, USA400 produced the least amount of biofilm as measured by crystal violet staining, yet showed relatively high CI values that were comparable to USA300. This discrepancy is perhaps driven by the fact that CI measurements are influenced by cell adherence, secretion of EPS, and cell spreading (44, 45), whereas crystal violet staining measures total biomass present including EPS, viable cells, and dead cells (46).

**Figure 1:**
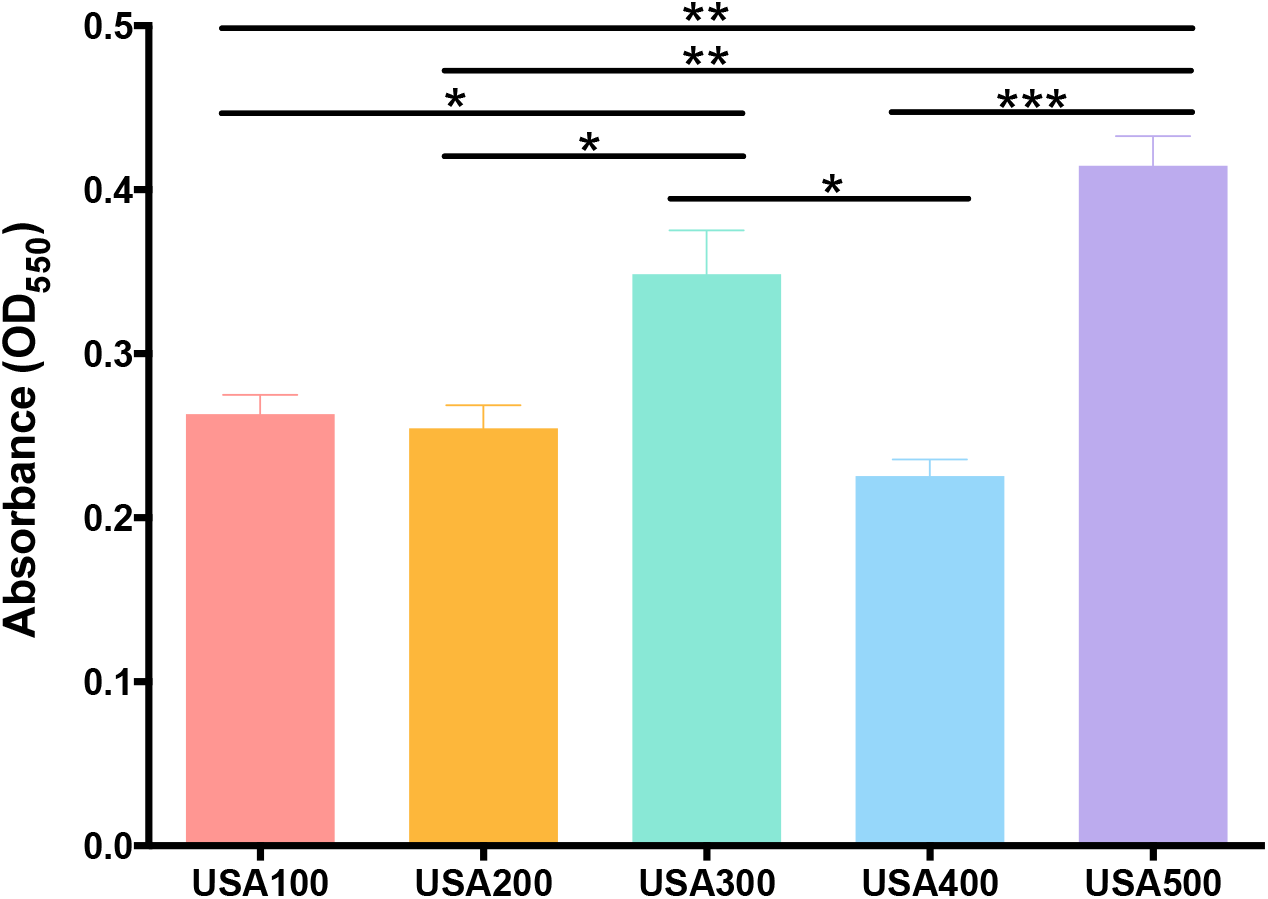
An assessment of biofilm formation across diverse MRSA clonal lineages. Static biofilms were grown in 96-well plates in TSB for 24 hours. Biofilm formation was quantified by crystal violet staining, followed by 100% ethanol elution and OD_550_ measurement. Assays were performed in biological triplicate with 8 technical replicates. Error bars represent ±SEM and Student’s *t-*test was used to determine statistical significance. *, P<0.05; **, P≤0.01; ***, P≤0.001.

**Figure 2:**
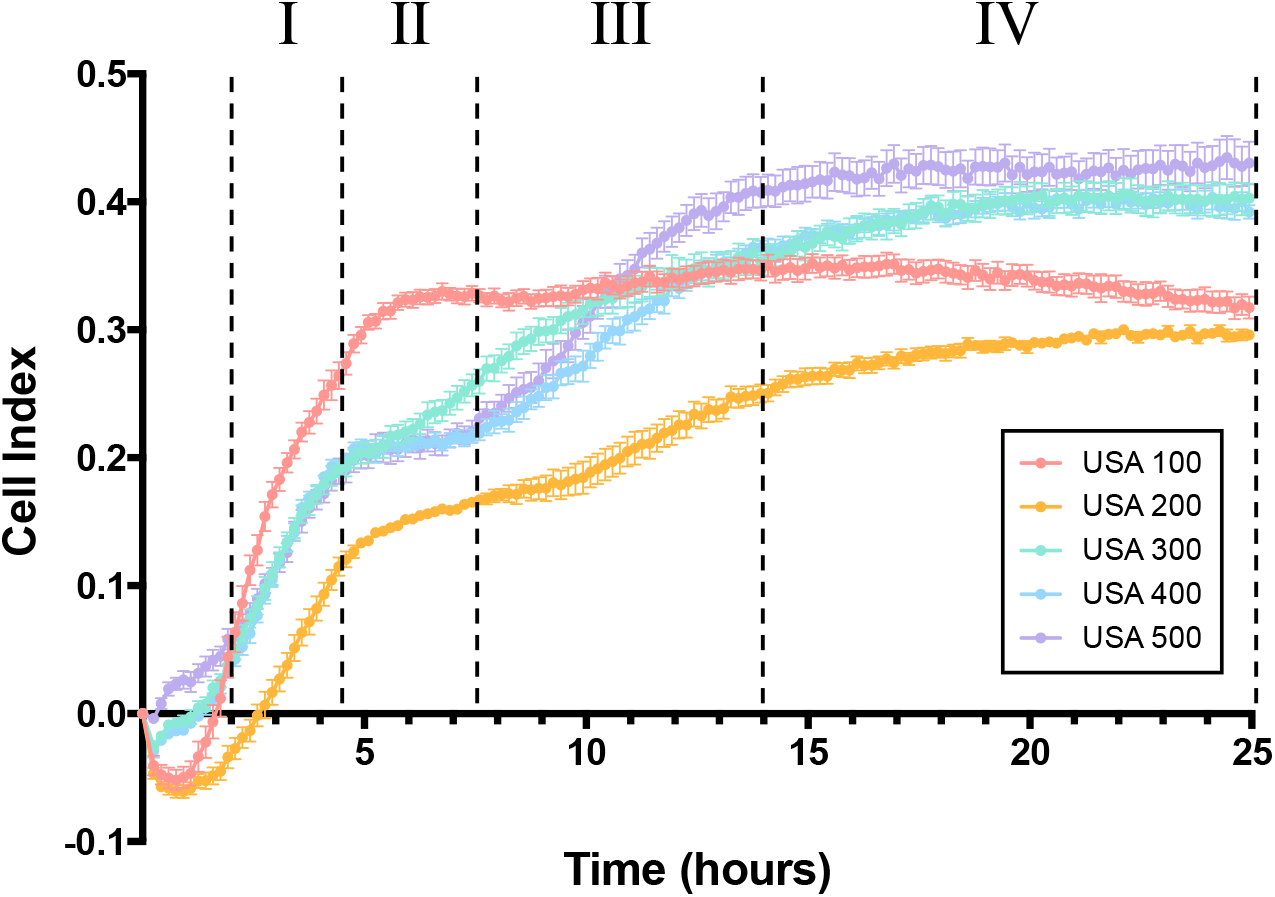
Real time analysis of *S. aureus* biofilm formation reveals distinct phases of development. Static biofilms were grown in specialized, electrode containing 96-well plates in TSB and monitored using a RTCA for 24 h. Each phase is separated by dashed lines and designated I, II, III, and IV. Phase I is characterized by a marked increase in CI (2 - 4.5 h) followed by a plateau during phase II (4.5 - 7.5 h). A second CI increase marks phase III (7.5 - 14 h) followed by levelling of CI values in phase IV (14 h onward). Data is presented from experiments repeated on three separate days with three biological replicates per strain in each instance. Error bars represent **±**SEM.

Beyond this comparison, our RTCA studies revealed four distinct phases of biofilm formation across the 5 strains. The first of these was characterized as a rapid phase of CI increase between 2 - 4.5 h, which we believe indicates attachment, with some strains (e.g. USA100) adhering more rapidly, whilst others (USA200) attaching more slowly. This phase was then followed by a plateau demonstrated by all strains, which lasted 3 hours for strains USA400 and USA500, although this was less apparent for strain USA300. Following this phase, biofilms other than USA100 then showed another CI increase, with USA500 being most accelerated. Finally, the CI values reach their maximum and remain steady for all strains, which signifies that the RTCA electrodes have been saturated and the biofilm has matured (44). Collectively, USA400 and USA500 very clearly exhibited these phases during analysis, whilst USA100 demonstrated a more prominent initial phase of attachment, quickly reaching its maximum cell index levels; with additional phases being less pronounced. As our data shows, each strain reaches a different maximum, with some achieving a CI greater than 0.4 (e.g. USA500) while others do not surpass 0.3 (USA200), which could be the result of variations in cell adherence, secretion of EPS, or cell spreading. The differences observed in biofilm formation and maturation are a testament to the diverse characteristics of these strains and the dynamic, evolving nature of biofilms.

### Differential expression in biofilm and planktonic cell populations drastically changes with population age

To gain more insight into the distinct phases observed during biofilm formation, and how biofilms evolve, we employed transcriptomic profiling of each isolate. RNA was collected from biofilm and planktonic cell populations after 5 h, 10 h, and 24 h of growth, corresponding to the end point of each phase observed during real-time tracking of biofilm formation, and subjected to RNA sequencing (Fig. 3, Fig. S1). Upon analysis, we observed that each clonal lineage exhibited distinct global expression profiles specific to its biofilm and planktonic growth (Table S1).

**Figure 3:**
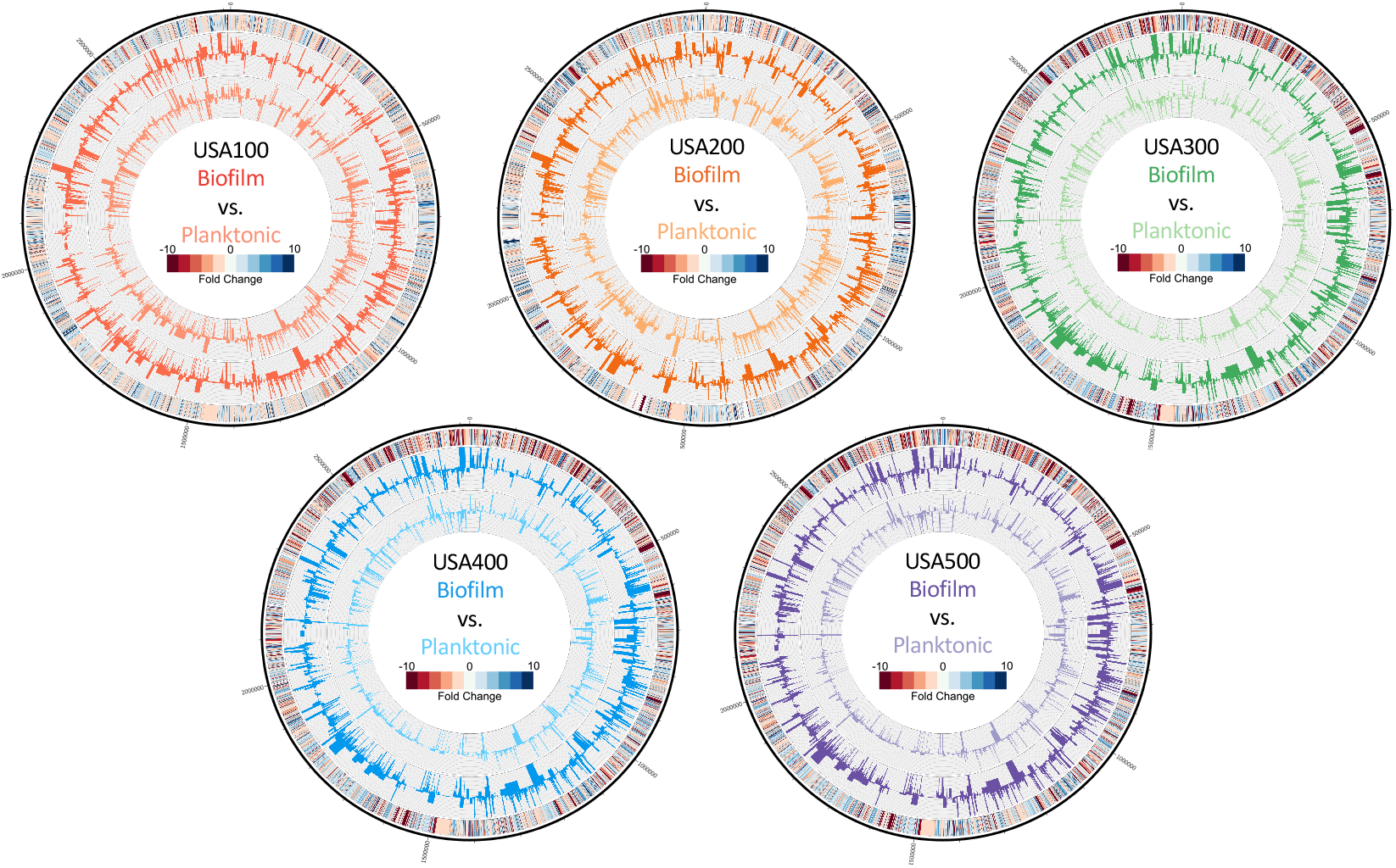
*S. aureus* biofilms exhibit differential expression compared to planktonic cell populations. Genomic maps were created for each strain depicting changes in the planktonic (inner histograms, light colors) and biofilm (outer histograms, dark colors) transcriptomes at 5 h reported as TPM expression values. The outermost circle is a heat map demonstrating fold change in expression, where red or blue indicates higher expression in the biofilm or planktonic cell population, respectively.

To explore this data more fully, we first chose to identify conserved gene expression patterns amongst all strains. To identify homologous genes, we used reciprocal BLASTn (Table S2) and identified genes with ≥ 3-fold differential expression between biofilm and planktonic cell populations. It is noted that this approach has its limitations, as only homologous genes are considered during this comparison, and strain specific genomic architecture is excluded; however, it does allow for a broad comparison of data between isolates. Genes showing preferential expression in biofilm and planktonic populations were then compared between strains to identify conserved expression patterns (Fig. 4). Preferential gene expression within biofilm and planktonic populations was largely strain dependent, with few genes showing ≥ 3-fold upregulation in all biofilms.

**Figure 4:**
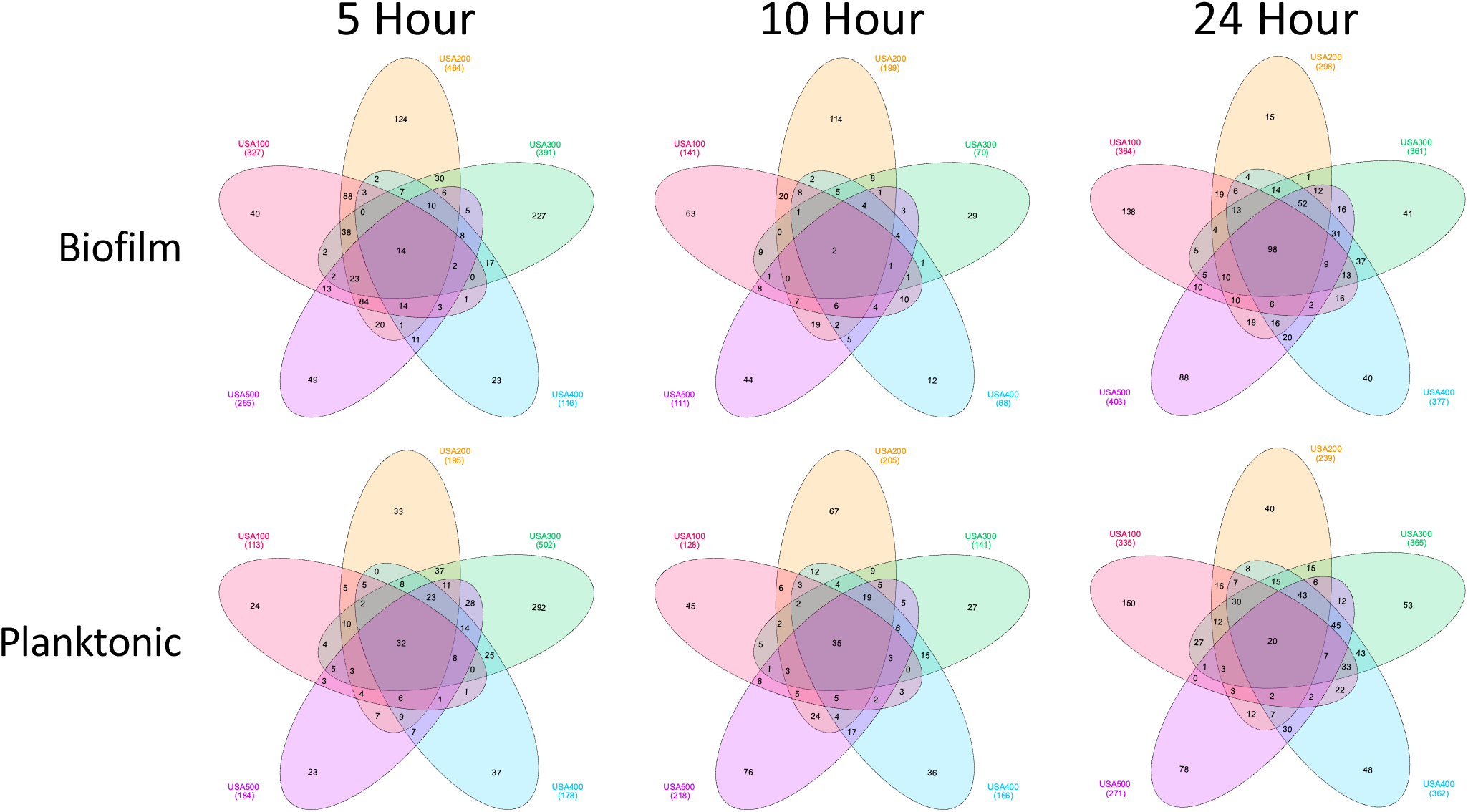
Preferential gene expression within biofilm and planktonic populations is largely strain dependent. Venn diagrams comparing biofilm and planktonic associated gene expression. Homologous genes with > 3-fold preferential expression in biofilm (top) versus planktonic (bottom) populations are shown for each timepoint sampled.

The majority of genes showing similar expression in all strains were preferentially expressed in planktonic cell populations at 5 h and 10 h, whereas at 24 h, the majority were preferentially expressed in biofilms (Fig. 5A, Table S3). Additionally, an interesting trend was observed regarding the number of genes expressed preferentially within biofilms. The number of common biofilm associated genes is low (n = 2) at 10 h and relatively high (n = 98) at 24 h. Compared to 5 h (n = 14), the number of common biofilm associated genes is 7-times greater at 24 h. When considering strains individually, each expressed a higher number of total biofilm associated genes at 5 h, which decreased by at least half for all strains at 10 h, and returned to comparable numbers at 24 h. Overall, this agrees with previous findings demonstrating cells within a biofilm have differential regulation compared to planktonic populations (47–49). Expanding on this model, we suggest the greater level of differential expressions within biofilms at 5 h and 24 h demonstrates that biofilm cells may be more physiologically diverse from their planktonic counterparts during attachment (5h) and maturation (24h) phases than during proliferation (10h). Furthermore, global transcriptional divergence between biofilm and planktonic cell populations occurred at 5 hours, indicating that differential regulation manifests earlier during biofilm development than previous works have considered. The majority of differentially expressed genes at this timepoint pertained to metabolic regulatory proteins (Table S3). It is possible that this evidence for reordered metabolism reflects metabolic dormancy of emerging persister cells, or is indicative of divergent metabolism in anaerobic regions forming within the biofilms (50, 51) (discussed below).

**Figure 5:**
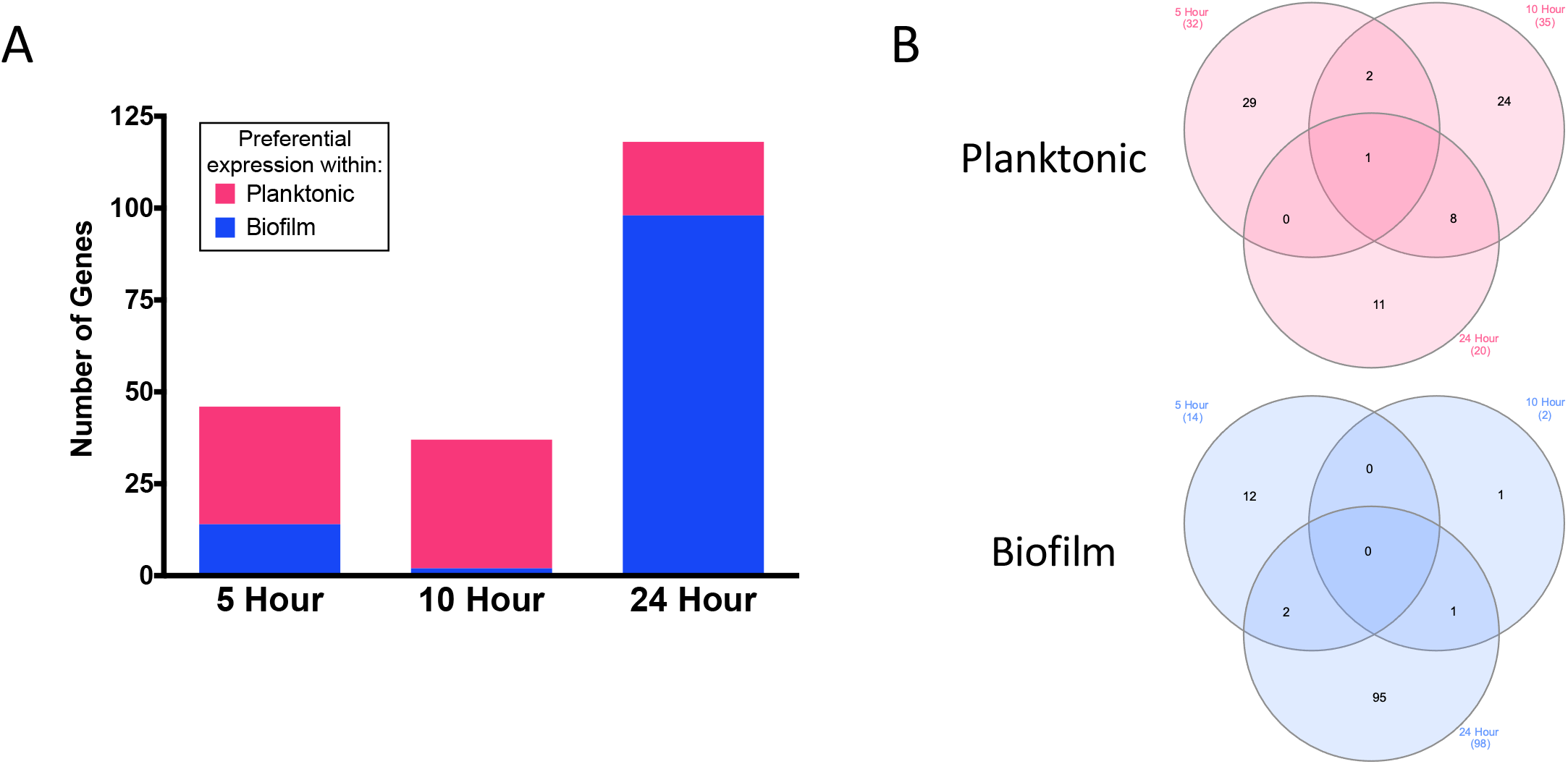
Differential gene expression occurs primarily in ageing biofilms. (A) The number of genes with > 3-fold preferential expression in biofilm or planktonic populations conserved across all strains at individual time points. (B) Venn diagram comparing biofilm and planktonic associated genes from (A) at each timepoint.

To validate our findings, a random selection of genes were assayed by RT-qPCR, revealing comparable changes to our RNA-seq data in all cases (Fig. S2).

### Urease production suggests acidification of the biofilm niche via fermentative metabolism

There was a near complete lack of constitutively expressed genes within biofilms, which reflects the dynamic, evolving nature of this population. Only a single gene showed preferential expression in one population over another for all strains (Fig. 5B, Table S3). This was *ureB*, which demonstrated lower expression in biofilm populations over planktonic cells, regardless of timepoint and strain. Further investigation of the full urease gene cluster (*ureABCEFGD*) revealed generally lower expression of each gene within biofilms. This operon encodes for the urease enzyme, which generates NH_3_ and CO_2_ from urea (52) and counteracts low pH caused by lactic acid, acetic acid, and formic acid accumulation (48). Previous studies have demonstrated a higher level of transcription of the urease operon in *S. aureus* biofilms (47, 48), however, this observation was made after 48 hours of growth, a timepoint beyond those sampled during our study. Therefore, upregulation of the urease gene cluster may occur only during prolonged growth in a biofilm state. In support of this, the absolute differential expression of the urease operon measured in our study decreased as time progressed, indicating that urease operon expression does gradually increase in biofilms (Fig. 6, Table S3).

**Figure 6:**
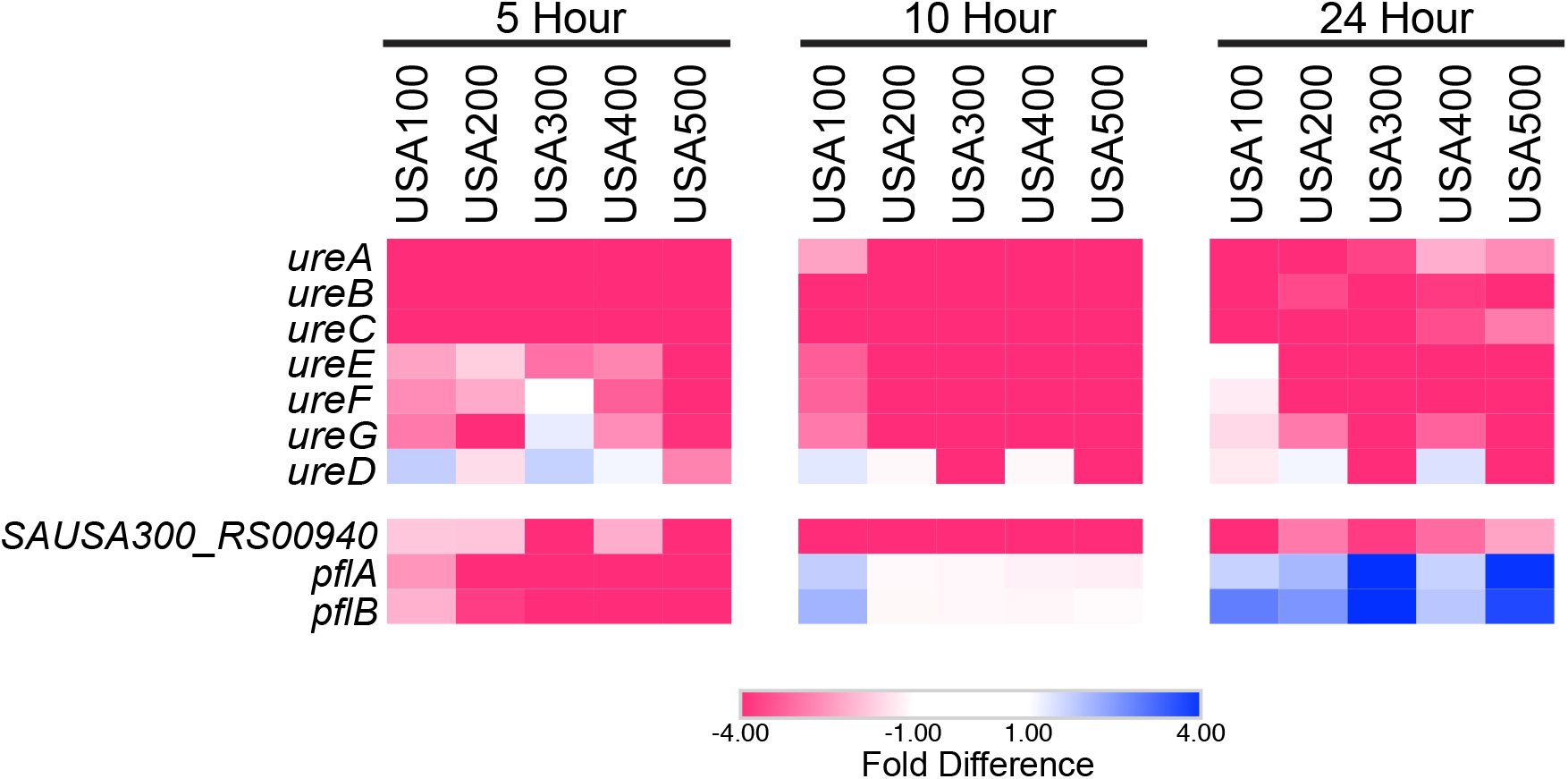
Biofilms show evidence of a formic acid metabolic response during later stages of growth. Shown are heatmaps depicting preferential gene expression in biofilm (blue) or planktonic (red) populations over time. Values for colors were assigned based on differences in expression between biofilm and planktonic cell population for each timepoint and each strain, all analyzed independently.

In agreement with Resch *et al.*, who observed a measurable increase in formate fermentation in biofilms, we saw evidence for the initiation of formate production in biofilms at 10 h. Specifically, we observed less transcription of NAD-dependent formate dehydrogenase (*SAUSA300_RS00940*) within 10 h biofilms (4.7 - 16-fold decrease, depending on the strain), which was less pronounced by 24 h (2.3 - 6.7-fold decrease) (Fig. 6). We also detected a gradual shift towards preferential expression of the genes encoding pyruvate formate-lyase, *pflA* (formate acetyltransferase-activating enzyme) and *pflB* (formate acetyltransferase). Under anoxic conditions within biofilms, PflA activates PflB to promote formate synthesis, which is then utilized for protein and purine production (53). Here, we noted a shift in *pflA* expression from 2.5 - 16.1-fold downregulation (5h), to 1.6 - 7.3-fold upregulation (24h) within biofilms. Similarly, transcription of *pflB* went from a 2.1 - 16.9-fold downregulation (5h) to 1.8 - 7.3-fold upregulation (24h) in biofilms. In agreement with our study, the work by Resch *et al.* detected *pflA* and *pflB* upregulation in biofilms by 16 h. Collectively, this implies that biofilms increase formate fermentation as they mature, and supports a model in which biofilms may counteract the accumulation of formic acid and consequential acidification of the biofilm by producing urease.

### Altered activity of stress response regulators speaks to the potential for their post translational regulation within biofilms

When looking specifically at 24 h data, we noted upregulation of numerous transcriptional regulators within biofilms, a phenomenon not observed for planktonic cells (Fig. 7). These included CtsR (SAUSA300_RS02715; ≥11.12-fold), LexA (≥3.58-fold), HrcA (≥6.78-fold), and SpxA (≥4.04-fold). CtsR, LexA, and HrcA all functionally act as repressors of various stress response elements. One would predict that their upregulation would lead to repression of their regulons, however, this does not appear to be the case. For example, CtsR represses the Clp family members *clpB, clpC, clpP (SAUSA300_RS04060)*, the *dnaK* operon, the *groESL* operon, and *ctsR* itself via direct promoter binding (54). Interestingly, all of these targets were strongly upregulated in 24 h biofilms (*clpB*: 5.02 - 120.28-fold; *clpC*: 1.53 - 15.75-fold; *clpP*: 4.35 - 14.58-fold; *dnaK*: 3.61 - 26.65-fold; *groES*: 4.50 - 40.44-fold; *groEL*: 3.05 - 28.81-fold). This is noteworthy because CtsR, SpxA, LexA, and HrcA are all subject to some form of post-translational regulation. Specifically, CtsR is phosphorylated by the co-transcribed protein, McsB, which marks it for degradation by the ClpCP protease complex under conditions of heat stress (55). HrcA requires the chaperonin, GroE, for proper folding; however, during heat stress, GroE becomes preoccupied refolding damaged proteins in a titrated fashion, leading to unresolved, improper folding of HrcA and de-repression of its heat shock regulon (56). SpxA acts as both a transcriptional repressor and activator, and is degraded by the ClpXP complex when bound by the YjbH protein. Conversely, under disulfide stress, SpxA abundance increases as YjbH-mediated ClpXP-catalyzed proteolysis decreases, leading to activation of the SpxA stress response regulon (57). In the presence of single-stranded DNA, RecA interacts with LexA, which triggers autocleavage of LexA (58). Importantly, a number of genes encoding these post-translational modifiers (*clpB, clpP, groES, groEL, recA, and yjbH*) are upregulated in the 24 h biofilm by ≥3-fold (Fig. 7). This suggests a mechanism by which the inactivation of CtsR, LexA, and HrcA by post-translational modification possibly leads to enhanced expression of their respective stress response regulons in established biofilms. Considering the widely varying conditions within biofilms over time, the upregulation of an ensemble of stress-response regulons is perhaps expected. Indeed, previous studies have demonstrated a correlation between expression of the CstR, SpxA, LexA, and HrcA regulons and biofilm integrity (59–62), including the CstR-regulated Clp-proteases (63). Conversely, the inactivation of SpxA and its regulon by YjbH and Clp-proteases, has been previously shown to enhance *S. aureus* biofilm formation (60). This suggests that unlike CstR, LexA, and HrcA, deactivation of the SpxA regulon may be beneficial to biofilms. Collectively, it would appear logical for a multitude of stress response regulons to be activated in biofilms and, based on our findings, this is potentially occurring via post-translational deactivation of key stress response repressors in older biofilm populations.

**Figure 7:**
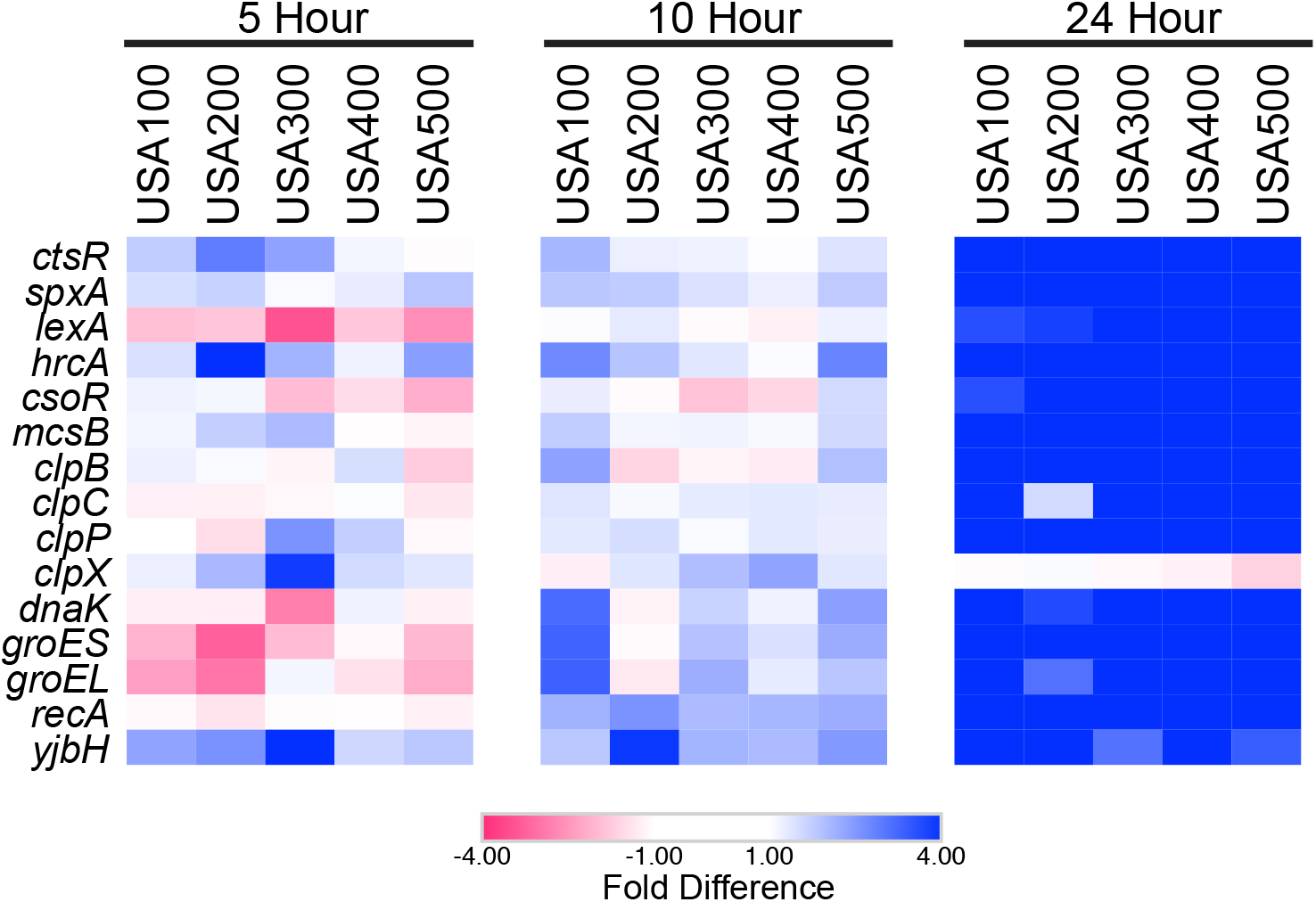
Transcriptional repressors and their post-translational modifiers are upregulated in 24 h biofilms. Shown are heatmaps that depict preferential gene expression in biofilm (blue) or planktonic (red) populations at 5 h, 10 h, and 24 h. Values for colors were assigned based on RNA-seq fold-difference in expression between biofilm and planktonic cell population for each timepoint and each strain, all analyzed independently.

### Developing biofilms display repression of factors modulating autolysis

When reviewing other specific changes in gene expression, we noted an interesting observation regarding *cidA* and *lrgA*. These genes encode a holin-like and antiholin-like murein hydrolase modulator, respectively, which antagonistically control cell lysis and genomic DNA release during biofilm development (64). Previous work suggests CidA activates murein hydrolases, which then triggers cell lysis, eDNA release, and biofilm adherence (64). LrgA has been shown to oligomerize with CidA and consequently antagonize CidA-mediated cell lysis (65). In 5 h biofilms, *lrgA* transcription was 3.88 - 13.10-fold higher, and *cidA* transcription was 4.31 - 12.98-fold lower (Fig. 8, Table S1). This suggests that at 5 h, biofilms might be less reliant on CidA-mediated eDNA release and adherence.

**Figure 8:**
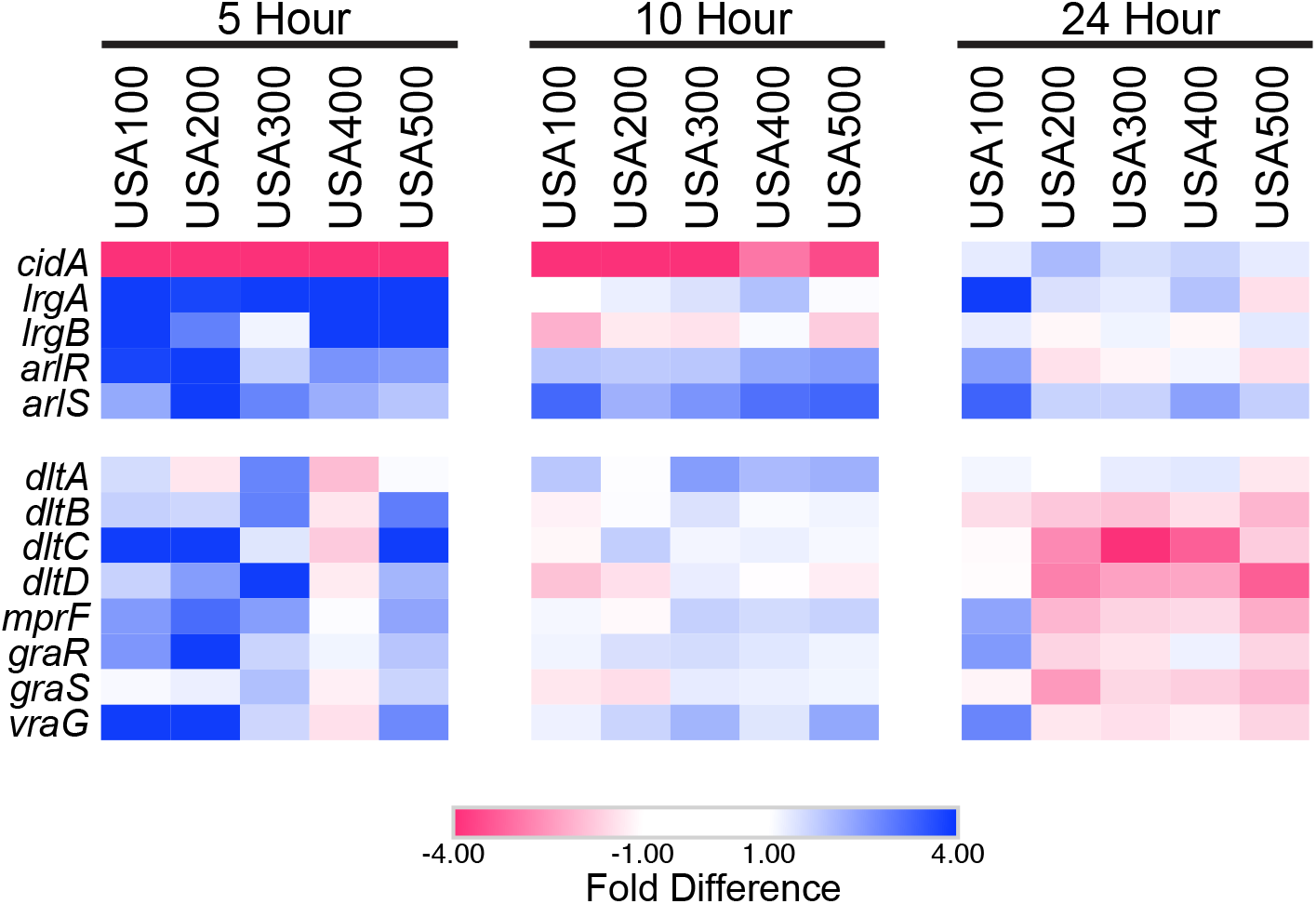
Initiating biofilms repress autolysis and activate PNAG regulators. Shown are heatmaps that depict preferential gene expression in biofilm (blue) or planktonic (red) populations at 5 h, 10 h, and 24 h. Values for colors were assigned based on RNA-seq fold-difference in expression between biofilm and planktonic cell population for each timepoint and each strain, all analyzed independently.

Coincidentally, phase I of biofilm formation is characterized by a rapid phase of attachment between 2 - 4.5 h, followed by a plateau in CI values starting at 5 h and continuing during phase II (Fig. 2). CI measurements are influenced by cell adherence and secretion of EPS (44, 45), and CidA-mediated cell lysis releases a sufficient amount of genomic DNA to mediate adherence during the initial stage of biofilm development (64). Therefore, decreased CidA production at 5 h leading to decreased eDNA incorporation into the biofilm matrix and decreased cell adherence may, at least in part, be responsible for the plateau of CI values observed at the end of phase I. In support of this model, inhibition of chemical lysis, and therefore eDNA release, by polyanethole sulfonate (PAS) has been shown to successfully inhibit biofilm formation, but only if PAS is added to the biofilm prior to hour 4 (64). This suggests that eDNA release is important for biofilm initiation, specifically prior to 4 h. Altogether, this supports the proposed model in which LrgA acts as an inhibitor of CidA, and our findings suggest that this antagonism may result in diminished reliance on CidA-mediated eDNA release at the end of phase I / beginning of phase II (5 h), prompting a shift in biofilm matrix development (Fig. 9).

**Figure 9:**
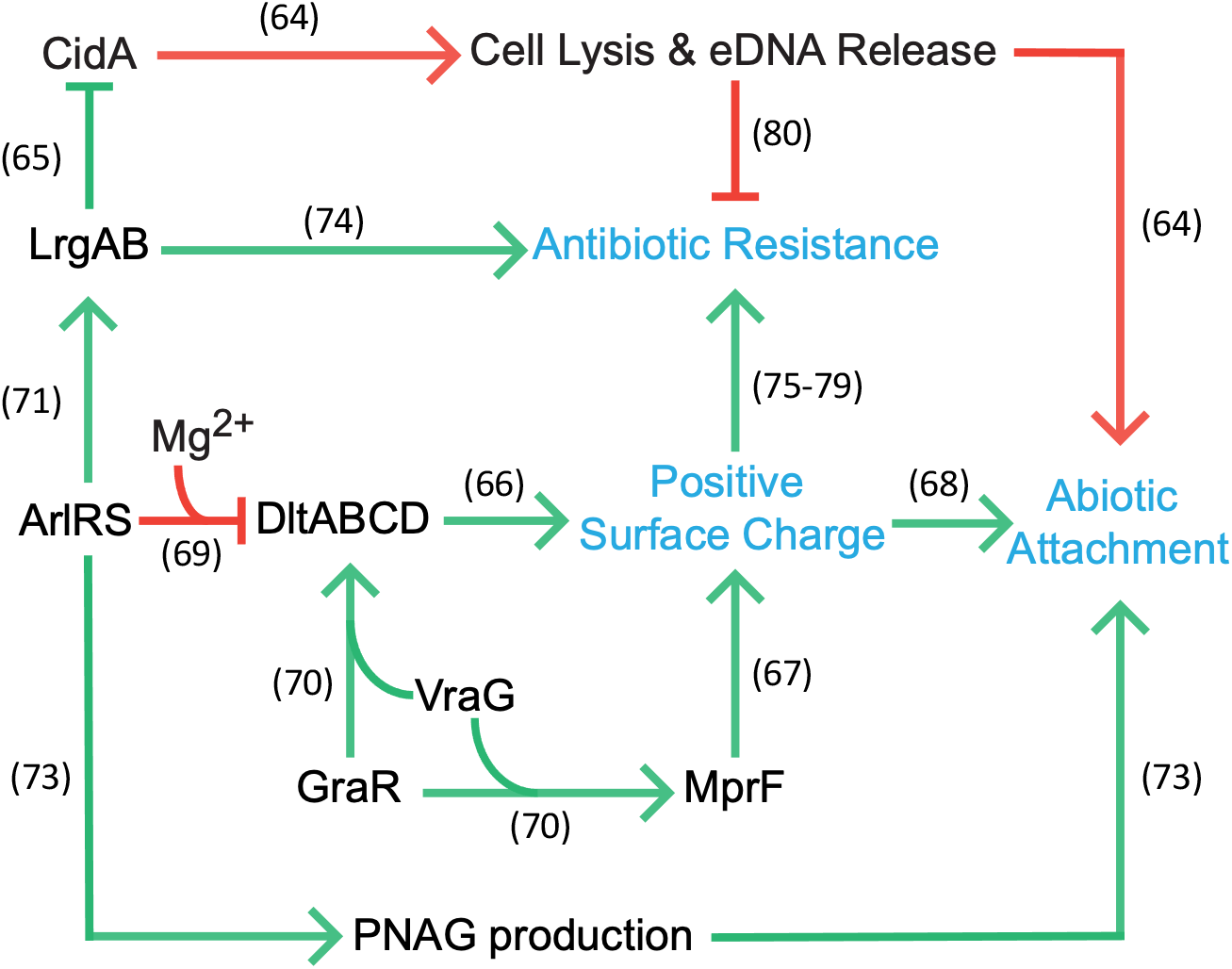
A proposed model for transcriptional regulation of biofilm-associated factors active after five hours of growth. The presented regulatory map depicts transcriptionally active (green) and inactive (red) pathways within 5 h biofilms. Points of regulation, and the resulting physiological changes which promote biofilm formation (blue), are based on previous studies (references shown).

### Cooperative regulation of positive cell surface charge and PNAG production during biofilm initiation

In addition to eDNA release, our data suggests that early biofilm development is potentially facilitated by positive cell surface charge. Two key determinants of net positive surface charge (*dltABCD* and *mprF*) were upregulated in biofilms at 5 h (Fig. 8). To increase positive surface charge, the products of the *dlt* operon (DltA, DltB, DltC, DltD) catalyze D-alanine incorporation into cell wall teichoic acids (66) and MprF modifies membrane phosphatidylglycerol with l-lysine (67). The electrical charge of *S. aureus* teichoic acids plays an important role in the initial steps of biofilm formation as *dlt* mutants have a net negative cell surface charge, and, despite wildtype levels of PNAG production, cannot colonize abiotic surfaces (68). In 5 h biofilms, the *dlt* operon and *mprF* (2.39 - 3.27-fold) showed greater expression in all strains, with the exception of USA400 (Fig. 8, Table S1). The ArlRS TCS has been demonstrated, in part, to repress *dlt* transcription and was upregulated during this phase (*arlR,* 1.91 - 4.73-fold; *arlS,* 1.71 - 8.27-fold). However, this particular regulatory cascade is dependent on the supplementation of media with cations, specifically Mg^2+^ (69) and therefore, is unlikely to be occurring under the conditions of our study. Alternatively, the expression of *mprF* and the *dlt* operon is also dependent on the co-transcription of response regulator *graR* of the TCS GraRS, and the downstream efflux pump *vraG* (70) (Fig. 9). Both *graR* and *vraG* were transcribed 1.65 - 7.69-fold and 1.61 - 6.94-fold higher, respectively, in biofilms compared to planktonic cells at 5 h. Again, the exception to this was USA400, which showed < 1.4-fold change for these genes (Fig. 8, Table S1). Considering that both *graR* and *vraG* are upregulated and the conditions for ArlRS regulation of *dlt* transcription are unfavorable in our study, our findings suggest that GraR and VraG influence *dlt* and *mprF* expression, possibly leading to increased positive surface charge and subsequent enhanced abiotic surface attachment for biofilms at 5h.

Another consideration is that, although repression of *dlt* transcription by ArlRS may not be occurring under our experimental conditions, ArlRS could be fulfilling other roles - promoting *lrgAB* expression (71), repressing autolysis (72), and/or promoting PNAG production and biofilm attachment (73). Indeed, enhanced transcription of *arlR* and *arlS* in biofilm populations (Fig. 8, Table S1) would allude to increased PNAG production in 5 h biofilms, which is consistent with our model. Collectively, this supports a scenario in which 5 h biofilms may employ PNAG-mediated attachment and a positive cell surface charge in order to promote abiotic attachment (Fig. 9).

It is also important to note that many of the above factors expressed in 5 h biofilms actively promote antibiotic resistance, whereas those that showed reduced expression hamper resistance. For example, LrgAB, MprF, VraR, and ArlRS promote beta-lactam tolerance (74–77) and were upregulated. Similarly, GraRS and DltABCD have been shown to promote resistance to vancomycin and other glycopeptides (78, 79) and their expression was also upregulated. Conversely, CidA abrogates beta-lactam resistance (80) and was downregulated. We therefore suspect that the cooperative effort of this regulatory network may be responsible, at least in part, for the increased tolerance of biofilms to antibiotics.

### Global pathway analysis reveals decreased production of translational machinery and ideal conditions for persister cell formation in early biofilms

Gene-level expression comparisons provide useful insight; however, pathway-level analysis offers a holistic view of how transcriptional changes are impacting the cell. For this approach, we first organized each gene into hierarchical functional clusters based on KEGG pathway ontology (81). Using the resulting clusters, a heatmap was generated from values of fold-difference in expression between biofilm and planktonic cells (Fig. S3). This metric revealed reduced transcription of the majority of the protein synthesis machinery in biofilms after 10 hours of growth (Fig. 10). Of these, the transcription of ribosomal protein genes *rplX, rplE, rpsN, rpsH, rplF,* and *rplR,* which are co-transcribed, demonstrated ≤ 3-fold decrease in expression in all biofilm cell populations at 10 h compared to planktonic cells. These genes encode 50S (Rpl) and 30S (Rps) subunit proteins, which facilitate proper subunit assembly. During assembly, ribosomal proteins S14 (RpsN) and S8 (RpsH) bind the 16S rRNA and coordinate assembly of the 30S subunit (82). Ribosomal proteins L24 (RplX) and L6 (RplF) bind the 23S rRNA, whilst directing 50S subunit assembly, and stabilizing 23S rRNA secondary structure (82). Ribosomal proteins L18 (RplR) and L5 (RplE) bind the 5S rRNA and are thought to mediate its attachment to the 50S subunit (82). Decreased production of these six key ribosome assembly proteins, and indeed an overall 53% of ribosomal proteins at 10 h across all strains, suggests that biofilms have diminished translational capacity. Impaired translational function is a hallmark of physiological dormancy – a condition which fosters persister cell formation (83). It is well known that persister cells form much more frequently within biofilms compared to planktonic populations and that their altered physiological behavior contributes to antibiotic tolerance (84). To our knowledge, it is not yet known how soon persister cells form, but it would appear that diminished translational capacity may contribute to this, and that they are likely present within biofilm populations by 10 h of growth.

**Figure 10:**
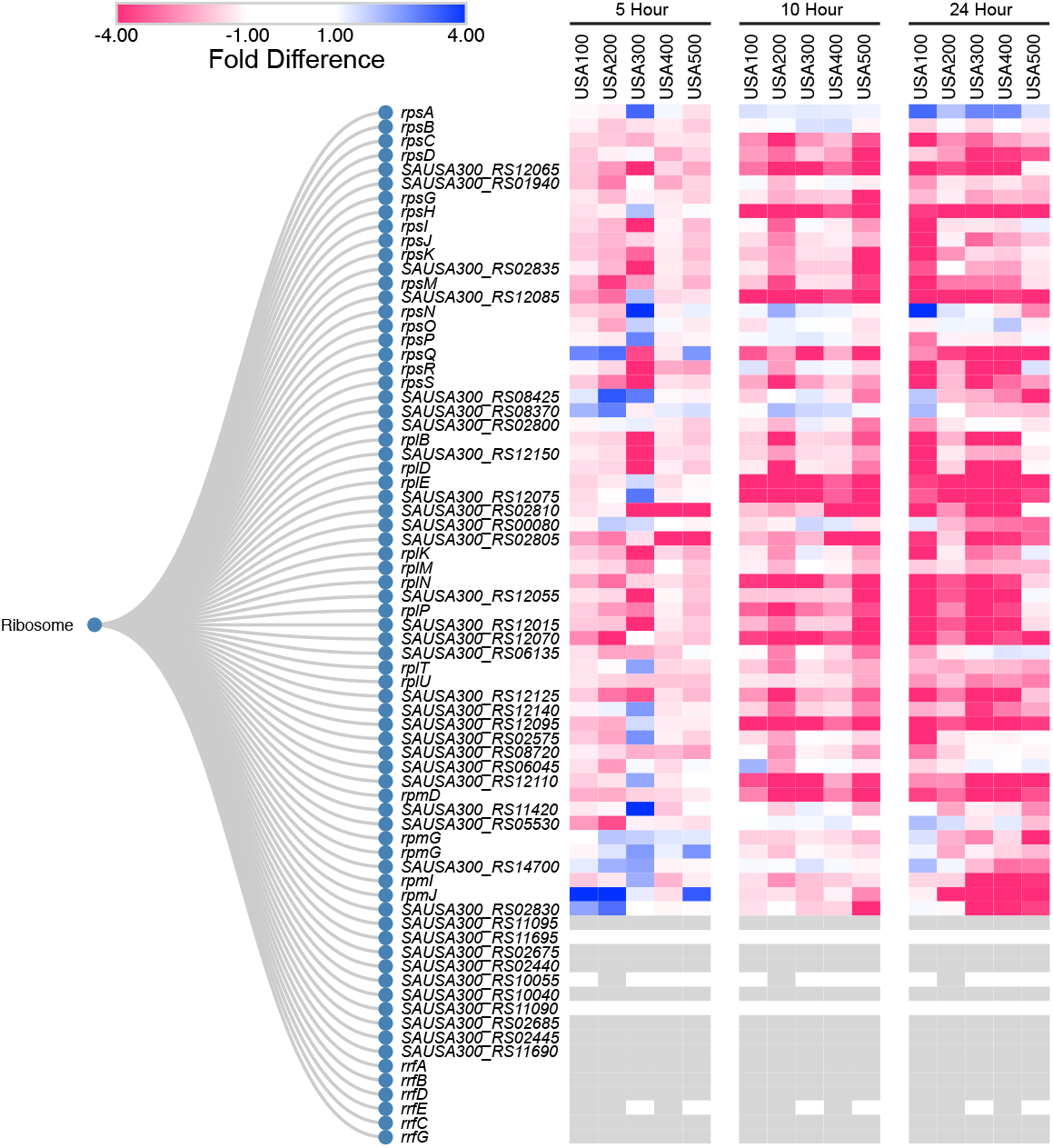
Pathway-level analysis reveals decreased production of the translational machinery within biofilms. Listed are homologous ribosomal genes organized by KEGG ontological function. USA300 locus tags are used throughout. The heatmaps depict levels of preferential expression in biofilms (blue) or planktonic (red) populations at 5 h, 10 h, and 24 h. Values for colors were assigned based on RNA-seq fold-difference in expression between biofilm and planktonic cell population for each timepoint and each strain analyzed independently.

### Biofilms display a shift towards nitrogen metabolism and anaerobic respiration

In addition to evidence of physiological dormancy, transcription within biofilm cells also reflects an altered metabolic state at later stages of growth. In particular, planktonic cells showed preference for *atpABCDEFGH* (the ATP synthase) expression at 24 hours, which is a major source of energy generation during aerobic respiration (Fig. 11). Preferred expression of ATP synthase within planktonic cells, and therefore lower expression within biofilms, suggests that mature biofilms employ alternative energy generation pathways that may mirror anaerobiosis (85). Indeed, biofilms develop anaerobic regions (50, 51), and anaerobic metabolism has been recognized as the preferred metabolic process for bacteria within deeper portions of fully formed biofilms. Indeed, previous studies have detected increased expression of anaerobic pathways (47, 49) within biofilms and anaerobic conditions stimulate greater biofilm formation compared to aerobic conditions (86). Our work shows that gene expression patterns within mature (24 h) biofilms skew significantly towards factors involved in an anaerobic state of respiration. Under anaerobic growth conditions, it has been shown that *S. aureus* increases the transcription if its alcohol dehydrogenases (*adhE, adh*) to regenerate NAD, and, even in the absence of nitrate, increases the expression of nitrate respiration (*narHIJ*) and nitrate reduction (*nirD*) genes (87). Each of the *S. aureus* biofilms in this study demonstrated enhanced transcription for all of these genes after 24 hours of growth, although it was to a lesser extent in USA400 (Fig. 11, Table S1). These findings are in addition to the three proteins associated with fermentative metabolism discussed above, *SAUSA300_RS9402, plfA,* and *plfB*, which showed increasing expression within biofilm populations as time progressed. Collectively, this supports a model in which biofilms possibly favor anaerobic respiration, with our data suggesting that this shift occurs during later growth phases. Interestingly, anaerobic growth and the repression of ATP synthase has also been demonstrated to enhance antimicrobial resistance in *S. aureus* (88, 89); and reduced levels of intracellular ATP is a hallmark-phenotype of *S. aureus* persister cell formation (85).

**Figure 11:**
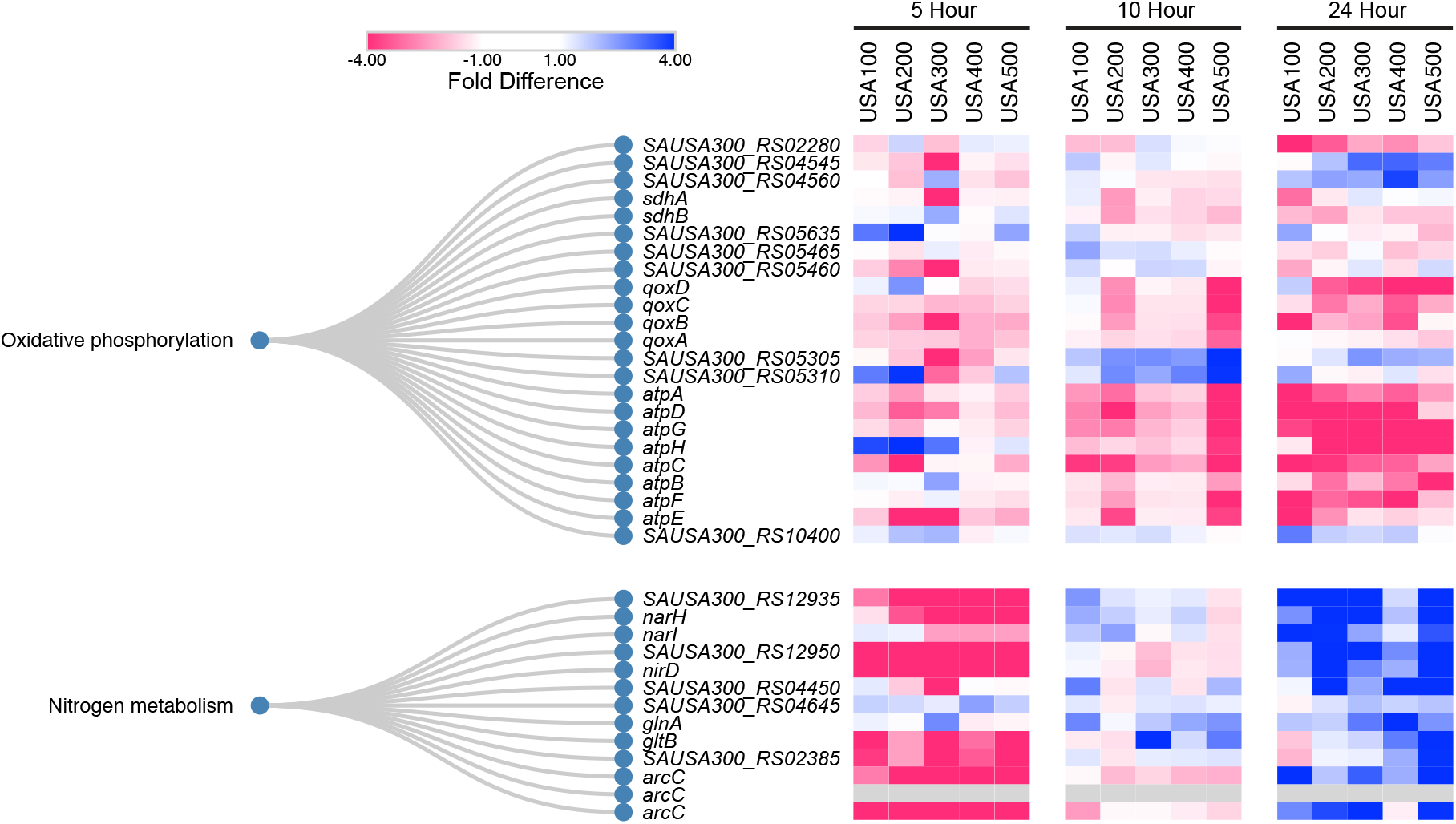
Mature biofilms increase nitrogen metabolism and reduce ATP synthesis. Listed are homologous genes organized by KEGG ontological functions, pertaining to oxidative phosphorylation and nitrogen metabolism. USA300 locus tags are used throughout. The heatmaps depicts levels of preferential expression in biofilms (blue) or planktonic (red) populations at 5 h, 10 h, and 24 h. Values for colors were assigned based on RNA-seq fold-difference in expression between biofilm and planktonic cell population for each timepoint and each strain analyzed independently.

### Differential expression of adherence factors provides insight into strain-specific infectious niche specialization

Unlike planktonic cells, as biofilms develop, various attachment factors that facilitate cell-cell adherence and attachment to extracellular matrix proteins generally increase in expression. When comparing the expression of known biotic surface attachment factors, including those that induce cell-to-cell adherence, several were not expressed within biofilms until 24 h. For example, the fibrinogen binding protein, Clumping Factor A (*clfA*) (13), showed preferential transcription in all 24 h biofilms, although to a lesser extent in USA100. ClfA was originally thought to be expressed early and facilitate initial attachment (90), however, our findings agree with recent work demonstrating stationary phase transcription of *clfA* (91, 92). Additionally, the exponential expression of *clfA* observed in previous work was during planktonic growth conditions supplemented with fibrinogen (90, 91). Thus, based on our findings, ClfA production likely does not occur until after biofilm formation has initiated, and initial attachment may not be dependent on ClfA. As discussed above, our results suggest that initial biofilm attachment is perhaps facilitated by CidA-mediated eDNA release followed by repression of this mechanism after 5 h by LrgA. At this time, biofilm attachment may be further enhanced by an increase in positive cell surface charge and enhanced PNAG production. Our data supports the notion that ClfA serves to maintain biofilm attachment at later time points, and perhaps provide late stage strengthening and maturation of the biofilm matrix.

As shown by tracking biofilm formation in real time, even closely related strains (e.g. USA300 and USA500) within a highly controlled environment present distinct phenotypes and trends (Fig. 2). This, in addition to variability in genomic architecture, suggests innate, idiosyncratic gene expression patterns exist within strains. As an example, preferential expression of *clfA* was less pronounced in USA100 biofilms (1.93-fold) compared to other 24 h biofilms (USA200, 13.25-fold; USA300, 18.83-fold; USA400, 13.49-fold; USA500, 61.30-fold). Like *clfA,* other genes encoding adherence factors, including Clumping Factor B (*clfB)*, IgG Binding Protein A (*spa)* (17), fibronectin-binding protein A (*fnbA*) and the Serine‐Aspartate Repeat Proteins (*sdrCDE*) (14), were preferentially expressed in USA300, USA400 and USA500 biofilms, whereas in USA100, these genes were more highly expressed in planktonic cells (Fig. 12). USA200 does not harbor the *sdrD* gene, but this strain also preferentially expressed *clfB, fnbA, sdrC, sdrE,* and to a lesser extent, *spa* in 24 h biofilms. Interestingly, overproduction or addition of soluble Spa in planktonic cultures can trigger bacterial aggregation in suspended culture (17). Such aggregates have enhanced cell-cell adherence, increased tolerance to antibiotics, are more resistant to shear stress, and are more mobile (93). Moreover, others have shown that the Sdr proteins facilitate platelet binding when cells experience shear stress in the upper range of arterial wall shear rates (91, 94), and that platelet binding is critical for *S. aureus* endocarditis infection (95). USA100 is the only strain to preferentially express these adherence factors in its planktonic cell population and is also the leading causes of bloodstream infections (20). Thus, it is entirely possible that the alteration of expression observed herein could contribute to this strains proclivity towards this type of infection.

**Figure 12:**
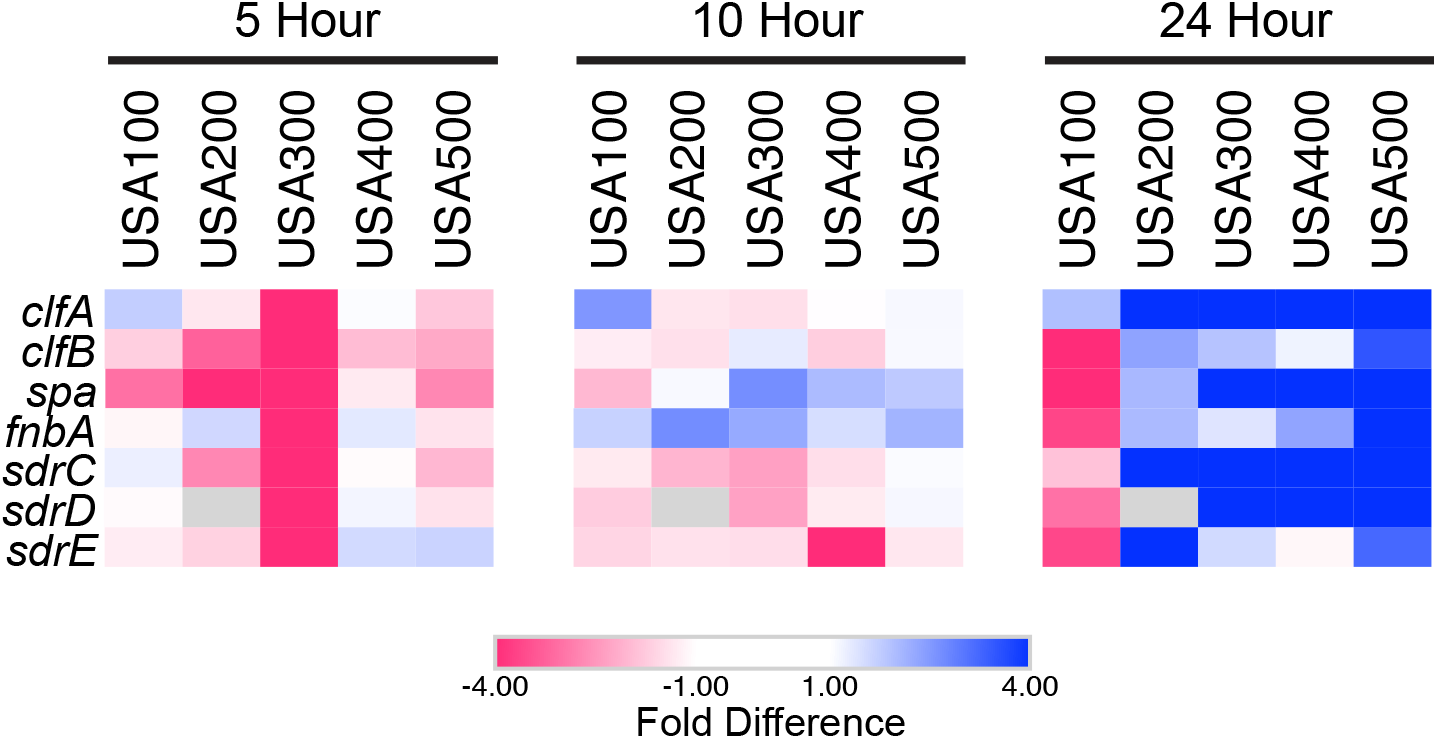
Strains display idiosyncratic expression of adherence and attachment factors. Shown are heatmaps that depict preferential gene expression in biofilm (blue) or planktonic (red) populations at 5 h, 10 h, and 24 h. Values for colors were assigned based on RNA-seq fold-difference in expression between biofilm and planktonic cell population for each timepoint and each strain, all analyzed independently.

## Concluding Remarks

Biofilms facilitate bacterial survival in diverse environmental niches, prolong infections, and continue to pose a major threat to patient recovery. *S. aureus* is capable of forming persistent biofilms, which are notoriously difficult to eradicate due to their increased tolerance to antibiotics and environmental stresses. Similar to their capacity for virulence, biofilm proficiency varies phenotypically amongst *S. aureus* clonal lineages. Much of what we currently know about biofilms is based on mature, preformed biofilm populations, therefore, little is known of the transient regulation driving attachment, proliferation, maturation, and dissemination. Our data herein highlights the diverse regulatory networks driving *S. aureus* biofilm formation. To our knowledge, this is the first study to investigate transcriptional regulation during the early, establishing phase of biofilms and compare global transcriptional regulation both temporally and across multiple clonal lineages. By monitoring biofilm formation of five *S. aureus* strains in real-time, we observe subtle differences in four distinct phases throughout biofilm initiation, proliferation, and maturation. The transcriptomic profiles, of both biofilm and planktonic cells, evolve drastically over time and exhibit differential expression between both populations throughout. We have uncovered a set of core transcriptional benchmarks, common to all clonal lineages, which includes a potential early shift toward anaerobic respiration in biofilms and reduced translational capacity. These findings are noteworthy because reduced cellular activity and an altered metabolic state have been previously shown to contribute to higher antibiotic tolerance and bacterial persistence. Furthermore, our data suggests that this shift may occur earlier than previously thought, prior to biofilms reaching maturity. Unfortunately, this also insinuates that the window of opportunity to avert biofilm infections via antibiotic therapy is likely short lived. Furthermore, the biofilms of all strains exhibited similar regulation of abiotic attachment factors, which highlights the importance of the ongoing efforts towards developing anti-biofilm surfaces or coatings for biomaterials and implants to prevent bacterial attachment. To this end, our findings support a model in which biofilms seemingly employ positive surface charge as a means for initial abiotic attachment, making the regulatory factors that control this process (VraG, GraR, MprF, and DltABCD) promising therapeutic targets to prevent biofilm initiation. Much of the strain-specific transcriptional regulation observed was that of well-studied virulence- and biotic attachment-factors. This is a testament to the unique regulatory strategy employed by each strain and gives insight into the factors behind their diverse infection styles. This study also provides a launching point towards understanding the highly orchestrated regulation driving each phase of biofilm development and will inform on future strategies to combat biofilm-mediated infections.

## Acknowledgements

We extend our thanks to the University of South Florida Genomics Program Genomics Equipment Core for the use of their facilities for RNA sequencing.

## Author contributions

Conceptualization: B.R.T., L.N.S.; investigation: B.R.T., M.E.M.; formal analysis: B.R.T.; writing – original draft preparation: B.R.T.; writing – review and editing: B.R.T., L.N.S.; Funding: L.N.S.

