## Supplemental Figure 1 for "A Global Transcriptomic Analysis of *Staphylococcus aureus* Biofilm Formation Across Diverse Clonal Lineages"

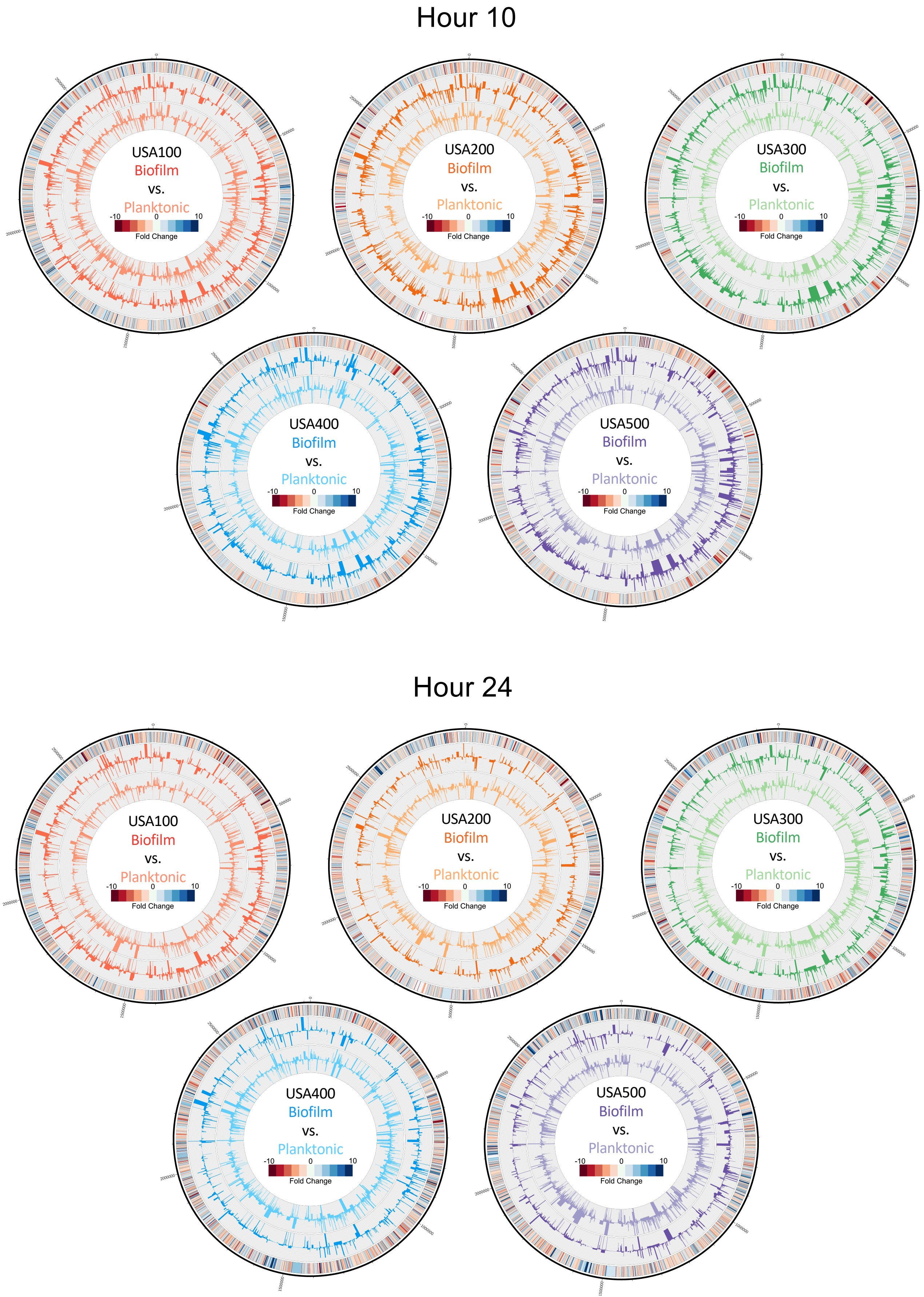


**Figure S1: *S. aureus* biofilms exhibit differential expression compared to planktonic cell populations.** Genomic maps were created for each strain depicting changes in the planktonic (inner histograms, light colors) and biofilm (outer histograms, dark colors) transcriptomes at 10 h (top) and 24 h (bottom) reported as TPM expression values. The outermost circle is a heat map demonstrating fold change in expression, where red or blue indicates higher expression in the biofilm or planktonic cell population, respectively.
