## Supplemental Figure 2 for "A Global Transcriptomic Analysis of *Staphylococcus aureus* Biofilm Formation Across Diverse Clonal Lineages"

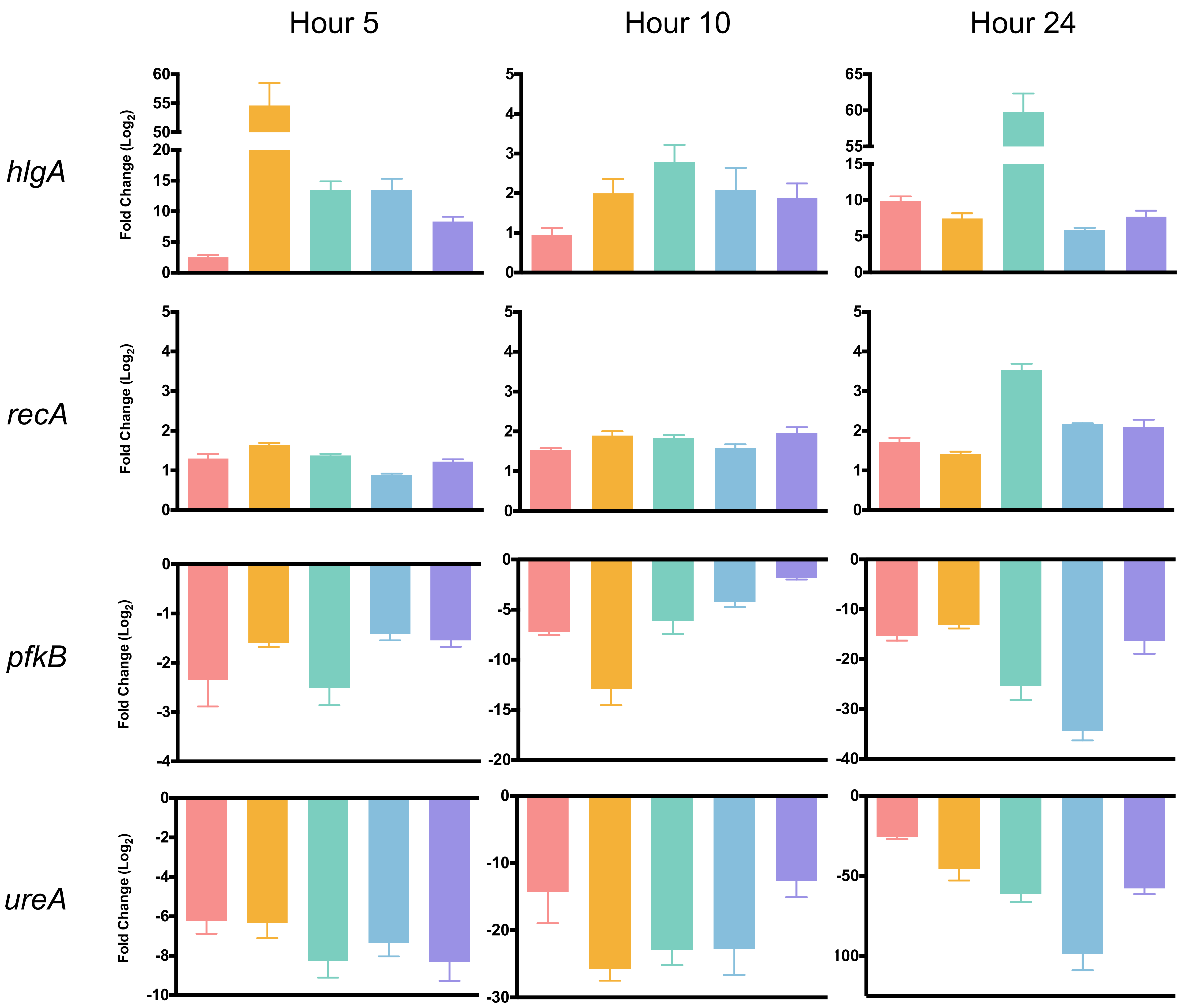


**Figure S2: RT-qPCR validation of RNA-seq findings.** RT-qPCR was performed using gene specific primers for four randomly selected genes (left), each timepoint (top), and each strain (USA100, red; USA200, orange; USA300, green; USA400, blue; USA500, purple). Expression levels were normalized to 16s rRNA and calculated using the $2^{-\Delta\Delta Ct}$ method. Data is reported as mean fold change of biofilm expression relative to planktonic expression ± standard error of the mean.
