## Supplemental Figure 3 for "A Global Transcriptomic Analysis of *Staphylococcus aureus* Biofilm Formation Across Diverse Clonal Lineages"

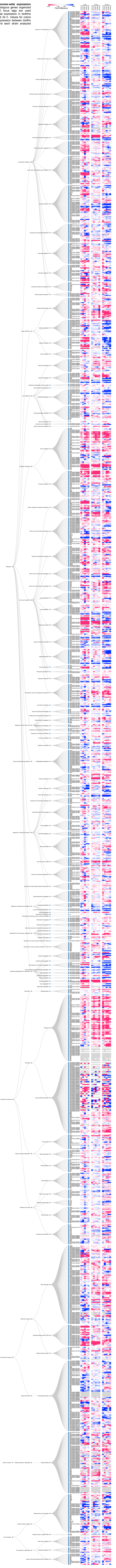

**Figure S3: Ontological dendrogram displaying genome-wide expression of *Staphylococcus aureus* biofilms.** Listed are homologous genes organized by KEGG ontological function hierarchies. USA300 locus tags are used throughout. The heatmaps depicts levels of preferential expression in biofilms (blue) or planktonic (red) populations at 5 h, 10 h, and 24 h. Values for colors were assigned based on RNA-seq fold-difference in expression between biofilm and planktonic cell population for each timepoint and each strain analyzed independently.
